# Understanding the dynamics of Telomerase-RNP complex in the human malaria parasite

**DOI:** 10.64898/2026.11.22.768470

**Authors:** Nirupa Chaurasia, Niharika Rai, Ariba Mahmood, Richa Shukla, Suchismita Bhowmik, Safia Baig, Neha Singh Chauhan, Nitin Mohan, Niti Kumar

**Affiliations:** Molecular Microbiology and Immunology Division, CSIR-Central Drug Research Institute, Sector 10, Jankipuram Extension, Lucknow, Uttar Pradesh-226031, India; Academy of Scientific and Innovative Research, Ghaziabad, Uttar Pradesh- 201002, India; Biological Sciences and Bioengineering, Indian Institute of Technology Kanpur, Uttar Pradesh 208016, India

**Keywords:** TERT, malaria, Telomerase-RNA, GAR1, NOP10, NHP2, Dyskerin, TCAB1

## Abstract

Amongst the human malaria causing species, *P. falciparum* exhibits an interesting telomere-telomerase (TERT) dynamics. In a striking contrast to humans, which have 10-15 kb long telomeres, parasite telomeres are shorter (1-2 kb) and are maintained by a very large *Pf* TERT (2.5 times bigger than hTERT) and its structural RNA (*Pf* TR is 5 times bigger than hTR). Till date, there are no studies on Telomerase-RNP complex in the malaria parasite. The RNP complex comprises of diverged TERT and RNA-interacting proteins (Dyskerin, GAR1, NHP2, NOP10, TCAB1) and structural TR. Interestingly, *Pf* TR undergoes stage-dependent transcription with non-uniform (heterogeneous) transcript length. The levels of *Pf* TR transcripts are much higher than that of *Pf* TERT, suggesting that *Pf* TR may have scaffolding non-coding RNA roles beyond TERT function. We also observed differential expression of *Pf* TERT and TR-interacting RNPs with non-stoichiometric cellular abundance. These RNPs can form higher-order complexes, which can be independent of TERT and TR. Through super-resolution microscopy (STORM), we observed distinct perinuclear localization of *Pf* TERT in ring, trophozoite and schizonts. Genotoxic and oxidative stress conditions affected the expression and subcellular fractionation of plasmodial Telomerase-RNP complex proteins (especially GAR1, NOP10), hinting towards their probable TERT-independent roles. Immunoprecipitation of Telomerase-RNP complex revealed specific enrichment of NOP10 and HSP90, hinting towards variable binding affinities of RNP proteins to *Pf* TERT. Our results suggest dynamics in the Telomerase-RNP assembly may allow them to perform both canonical and non-canonical functions to support intracellular parasite survival in evolutionary distinct hosts.

**Author Summary:** To support continuous growth, the cells have to maintain their telomeric ends during each round of DNA replication. Eukaryotes maintain their telomeric length and genome stability through specialized Telomerase-RNP complex that comprises of Telomerase reverse transcriptase (TERT), Telomerase RNA (TR) and its interacting proteins. TERT maintain telomeres by the addition of repetitive Guanine-rich sequences to the 3’-end of linear chromosomes to overcome the progressive DNA shortening due to end-replication problem. Malaria parasite has evolved a very unique Telomerase enzyme that maintains the telomeric ends and gives competitive survival advantage through continuous cellular growth throughout all the stages of its intracellular survival in the human and mosquito vector. Understanding the differences in the host and parasite Telomerase-RNP complex will open avenues for alternative sites of pharmacological intervention to tackle emerging drug resistance.

## Introduction

Malaria causing *Plasmodium* species exhibit a complex digenetic lifecycle involving multiple developmental stages in both the human host and female *Anopheles* mosquito vector (1). The parasite survives in different microenvironments, such as hepatocytes during the liver stage, erythrocytes in the blood stage in human, and midgut/salivary gland tissues in the mosquito. These transitions impose unique physiological challenges, including oxidative stress, immune threats, and rapid proliferation demands (2, 3). To ensure survival and successful transmission, this parasite must preserve its genome integrity across all stages of its lifecycle. *Plasmodium* has evolved specialized replication and repair systems for its nuclear and organellar (apicoplast and mitochondrion) genomes (4-7). The *P. falciparum* apicoplast genome is a ∼35 kb circular DNA that encodes rRNAs, tRNAs, ribosomal proteins, RNA polymerase subunits, and a limited set of housekeeping proteins. In contrast, the mitochondrial genome is a ∼6 kb linear concatemer that encodes three respiratory chain proteins (Cytb, Cox1, and Cox3) together with fragmented rRNAs (8).

The nucleus of *P. falciparum* harbors a 23 Mb AT-rich haploid genome distributed across 14 linear chromosomes. The chromosome ends have telomeres which are repetitive (GGGTTT/CA)_n_ sequences, approximately 1-2 kb in length. Telomeric ends are flanked by variable subtelomeric regions (30-120 kb) that encodes multicopy gene families (*var*, *rifin*, *stevor* genes), which mediate antigenic variation and immune evasion (9). Through Telomere position effect (TPE), *Plasmodium* telomeric structure can influence the transcription or heterochromatin state of subtelomeric regions coding for antigenic variation genes responsible for immune evasion (10-12). Unlike its human host, in which hTERT is only expressed in stem cells, germline and cancerous cells, the parasite expresses *Pf* TERT is expressed throughout the asexual and sexual stages of its lifecycle (13-15). This ensures that parasite telomere length is maintained during multiple rounds of schizogony.

Human Telomerase-RNP complex adopts bilobed architecture with one lobe having the catalytic core (comprising of TERT) and another H/ACA lobe containing hTR interacting proteins with a stoichiometric ratio of one molecule of TCAB1, and two molecules each of Dyskerin, GAR1, NHP2, NOP10. Cryo-EM structure also identified an additional transient interaction of H2A/H2B histone with the catalytic core (16-18). A recent study on hTR using chemical probing (DMS-MaPseq) and ensemble deconvolution reported the conformational heterogeneity and kinetic folding traps in hTR (19). In a striking contrast to human, *Pf* TERT is approximately 2.5 times larger than its hTERT, and its non-coding structural RNA (*Pf* TR, 2148 nt) is nearly five times longer than the hTR (451 nt). Due to the extremely large size and potential structural variability, domain organization of *Pf* TERT and *Pf* TR remains poorly understood (20, 21).

Till date, there are no studies on plasmodial telomerase-RNP complex, which limits our understanding of their role in assembly, trafficking, recruitment to telomeric ends, and adaptations to stress response for competitive survival in human host. In this work, we have tried to understand the evolutionary divergence and differential expression of the *Pf* Telomerase-RNP complex proteins (Dyskerin, GAR1, NHP2, NOP10, TCAB1). We observed stoichiometric imbalance with significantly higher transcript levels of TR and TR-interacting proteins over *Pf* TR, thereby suggesting their telomerase-independent roles and regulation. We also observed differential partitioning of Telomerase-RNP components especially for GAR1 and NOP10 suggesting their probable non-canonical roles under physiological and stress conditions. *Pf* TERT interacting partners also include HSPs, which may play an important role in TERT folding and assembly. Understanding the organization, assembly, and interacting partners of the parasite telomerase-RNP complex may provide valuable insights into its functional regulation and identify potential targets for selective pharmacological intervention.

## Results

### Evolutionary Divergence of Pf Telomerase-ribonucleocomplex

*Pf* TERT (catalytic core) is composed of 2518 amino acids and harbors long atypical insertions, which are not present in other TERTs (**Table S1, Figure S1**). This multidomain protein comprises of Telomerase Essential N-terminal (TEN) domain, the Telomerase RNA Binding Domain (TRBD), the Reverse Transcriptase (RT) domain, and the C-Terminal extension (CTE) (**Scheme 1**). The TEN domain plays role in capturing single-stranded telomeric DNA and interacts with the TR. Within the TRBD, conserved CP, QFP and T motifs contribute to the formation of the RNA-binding pocket. The RT domain is the enzymatic core of TERT consists of multiple motifs (1,2,A,B,C,D) (22). Interestingly, based on multialignments, *Pf* TERT domains do not share direct sequence similarity with human TERT, but they retain structural similarity in QFP motif in TRBD, B’,C,D motifs in RT that underpin their biological function **(Figure S2)**.

**Scheme 1:**
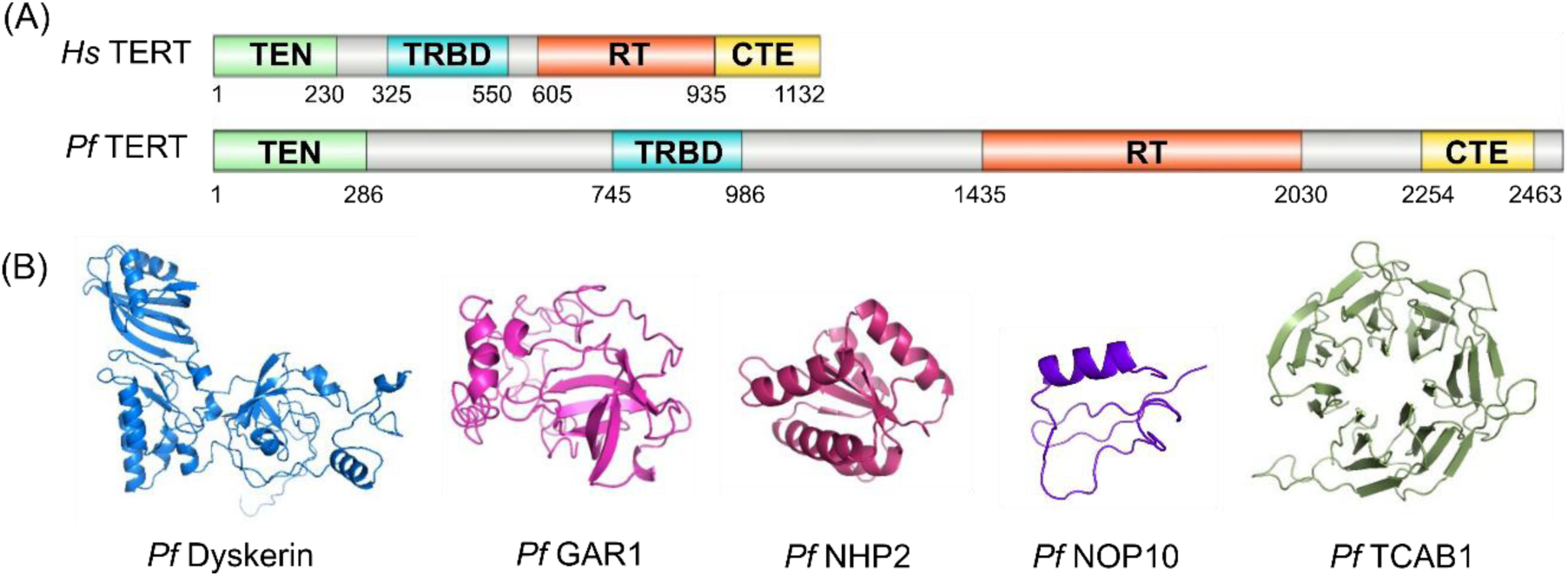
Human and *P. falciparum* Telomerase-RNP complex. (A) Domain organization of human and *P. falciparum* Telomerase reverse transcriptase (TERT). (B) Predicted models of plasmodial orthologs of Telomerase-RNP complex proteins using AlphaFold.

*Pf* TERT requires an exceptionally large non-coding structural RNA (*Pf* TR 2148 nt) in comparison to human TR (451 nt) to assemble into a functional Telomerase-RNP complex. Unlike the hTR, the *Pf* TR boundaries for Template, TBE, Pseudoknots, CR4/5 and H/ACA motifs are not well-defined (19). The 3′ end of the TR is stabilized by H/ACA binding dimeric proteins such as Dyskerin, Glycine-Arginine rich 1 (GAR1), Non-histone protein 2 (NHP2), Nuclear protein 10 (NOP10) (23, 24). Telomerase cajal box protein 1 (TCAB1) serves as an assembly factor for TERT recruitment on TR (25). The plasmodial TR-interacting proteins (Dyskerin, GAR1, NHP2, NOP10, TCAB1) harbor conserved structural folds despite low sequence similarity with human orthologs **(Table S2).**

### Expression Pf Telomerase-RNP complex proteins

Stage-specific protein expression showed that *Pf* TERT is expressed throughout all asexual blood stages, with moderately higher expression in trophozoite/schizont stages, wherein rapid nucleic acid synthesis occurs. Super-resolution (STORM) microscopy revealed distinct punctae-like signals of TERT near the nuclear periphery in ring, trophozoite and schizont stages (**Figure 1**). This distinct signal represents the *Pf* TERT recruitment to telomere-clusters anchored to the nuclear periphery, a phenomenon conserved across all eukaryotes (20).

**Figure 1:**
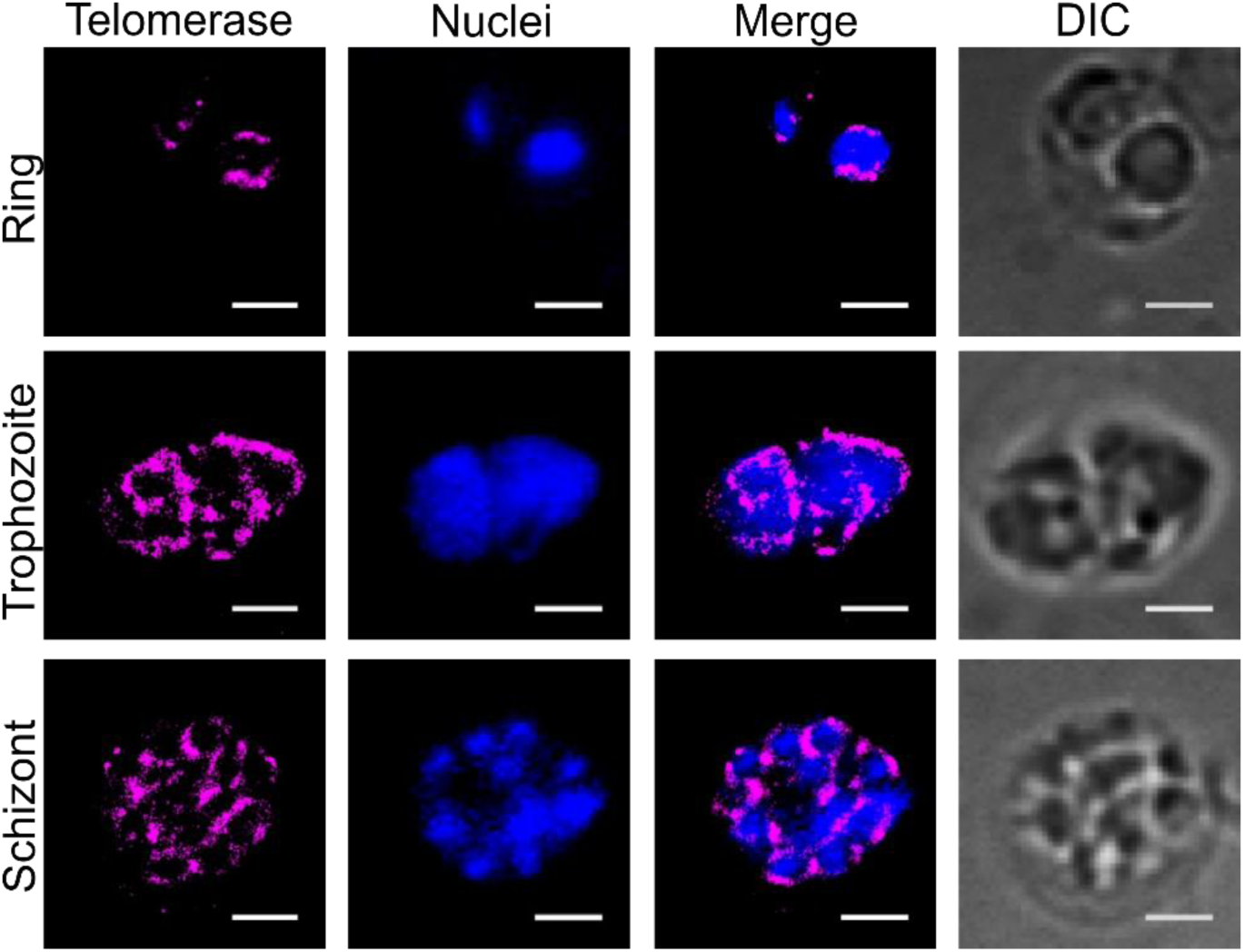
Stochastic Optical Reconstruction microscopy for localization of *Pf* Telomerase. Immunofluorescence was performed for PF3D7-infected RBCs at different stages of it’s lifecycle (ring, trophozoite and schizont). *Pf* Telomerase probed with AlexaFluor 647 and nuclei were stained with Sytox dye. Scale bar 2 µm.

*Pf* TERT assembles into an active RNP-complex through a 2148 nt long RNA (TR). However, we observed that *Pf* TR transcripts were significantly abundant than TERT transcripts across ring, trophozoite and schizont stages (**Figure S3**). Surprisingly, full length amplicon was not obtained through PCR from a reverse-transcribed template. Therefore, we checked the expression of *Pf* TR through PCR using sequence-specific primers spanning from the 5’-3’ end of the transcript (**Table S3**). The genomic-DNA (gDNA) showed amplicons of desired length for all primer sets (**Figure 2**). Upon PCR from reverse transcribed template, the amplification with different gene-specific primers was not very consistent. Further, the amplicon intensity was different across ring, trophozoite and schizont stages (**Figure 2, Figure S4A**). The plausible explanation for this observation can be (a) stage-specific expression of *Pf* TR and post-transcriptional modifications, (b) the existence of transcripts of variable lengths of *Pf* TR and/or (c) synthesis of cDNA with internal deletions due to slippage of reverse-transcriptase enzyme as it encounters stem/loop secondary structures (due to polyA-polyT stretches) in *Pf* TR. The available RNAseq data (PlasmoDB, **Figure S4B**) also show a similar profile of low transcriptional activity across specific region of transcript length which is in agreement with the above observations.

**Figure 2:**
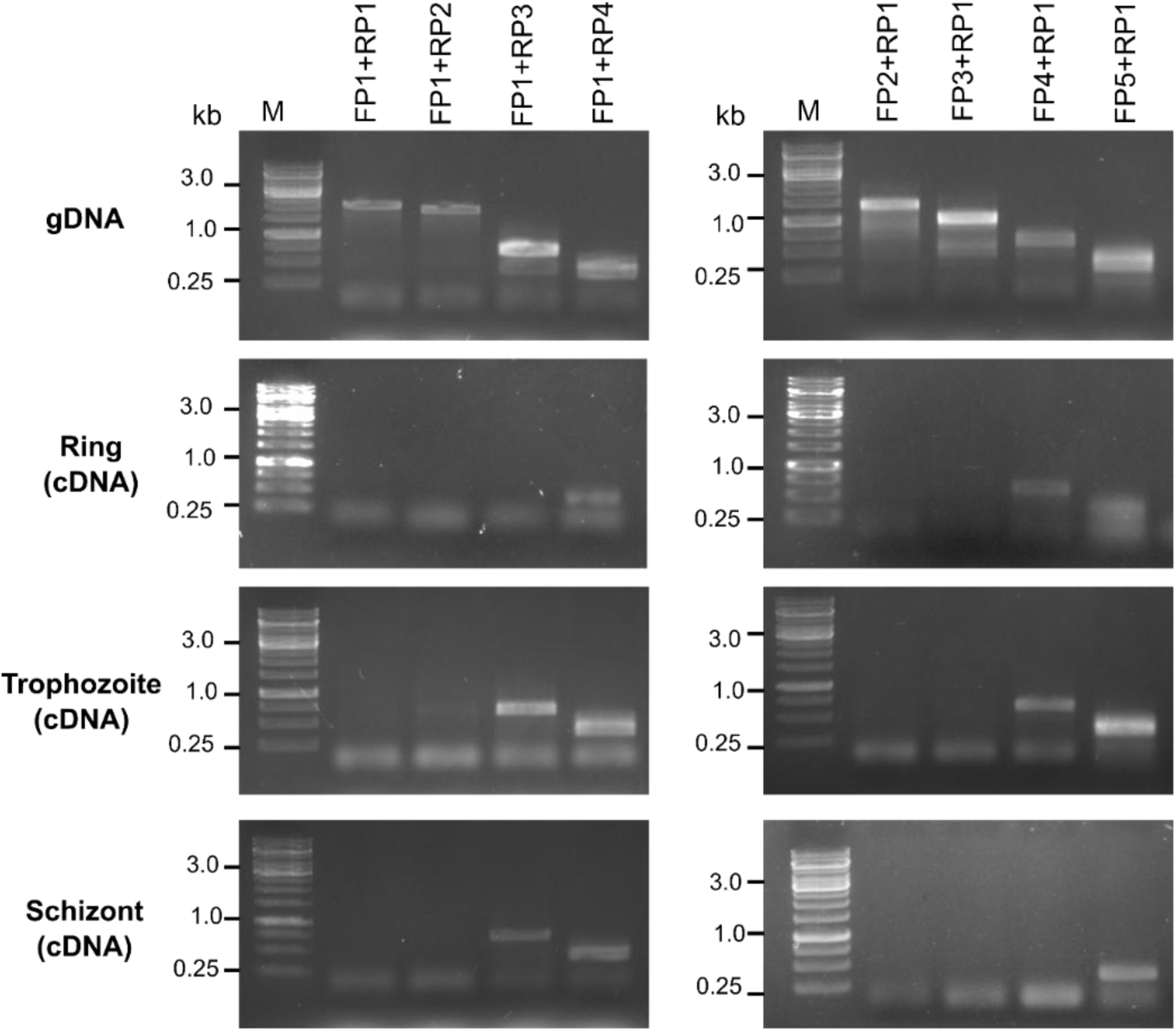
*Pf* Telomerase RNA (TR) amplification at different asexual stages. Using specific PCR primer-set covering the TR length, PCR amplication was performed using the gDNA and cDNA (prepared from ring, trophozoite and schizont stages).

We checked the cellular expression of other proteins interacting with *Pf* TR that form the Telomerase-RNP complex. We observed that transcript levels of TR-interacting proteins (GAR1, Dyskerin, NHP2, NOP10, TCAB1) were higher than TERT (Fig S5). The differences in the stage-specific expression suggests that these may have role beyond *Pf* Telomerase-RNP complex. NHP2 and NOP10 had similar expression levels throughout ring, trophozoite and schizont stages. Dyskerin and GAR1 exhibited very low expression at the ring stage, in comparison to their respective trophozoite and schizont stages. TCAB1 shows visible protein expression only in the ring stage (**Figure 3A**). To examine the effect of genotoxic and oxidative stress on the expression of these proteins, the parasite was exposed to 0.025% MMS, a DNA-damaging agent, or 7.5 µM FCCP, a mitochondrial uncoupler during trophozoite stages (30-32 hpi, for 6 h). Giemsa stained microscopy images showed healthy trophozoites in control culture, but MMS and FCCP treated culture had compact trophozoites. Under these stress conditions, expression of telomerase and its associated H/ACA proteins was downregulated, while heat shock proteins (HSP70 and HSP90) were upregulated **(Figure 3B).**

**Figure 3:**
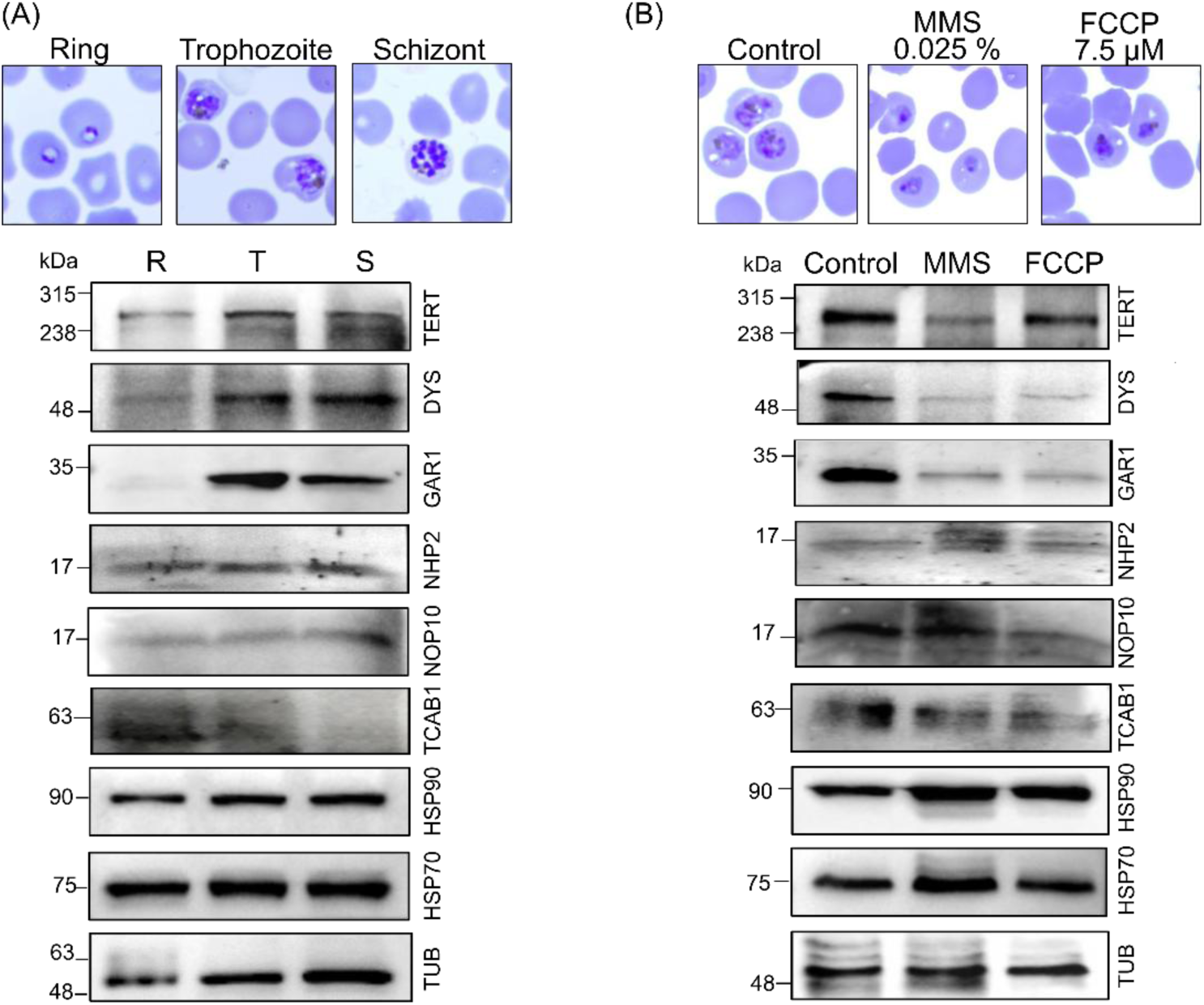
Expression of Telomerase-RNP complex under physiological and stress conditions. (A) Giemsa-stained images of PF3D7 for ring, trophozoite and schizont stage with corresponding western blots for cellular expression of *Pf* Telomerase-RNP components using the respective sera. (B) Giemsa images showing phenotypic changes upon exposure to 0.025 % MMS or 7.5 µM FCCP at trophozoite stage (30 hpi) for 5.5 h and corresponding western blots for cellular expression of Telomerase-RNP components upon exposure to stress.

### Cellular fractionation of Pf Telomerase-RNP complex proteins

Subcellular localization studies under physiological conditions showed that *Pf* TERT is predominantly present in the nuclear compartment (**Figure 4**). Telomerase-RNP complex proteins like, Dyskerin, NHP2, and NOP10 localize predominantly in the nucleus, while GAR1 is distributed in both nuclear and cytoplasmic compartments. As the expression of TCAB1 is very low in the trophozoite stage (**Figure 3A**), the fractionation data shows faint signal in both nuclear and cytoplasmic fractions. In contrast, HSPs are primarily cytoplasmic, histone and tubulin served as the respective control for nuclear and cytoplasmic compartments. Upon DNA damage or mitochondrial stress, we observed differential localization of components of the Telomerase-RNP complex. *Pf* TERT and Dyskerin partitioned in both nuclear and cytoplasmic fractions, but NHP2 localization did not change upon DNA or mitochondrial stress. Interestingly, GAR1 showed predominant cytoplasmic localization specifically during DNA damage and NOP10 predominantly localized in cytoplasm during mitochondrial stress. HSPs showed stress-dependent differential localization specifically during mitochondrial stress with both HSP70 and HSP90 partitioning into the nuclear fraction (**Figure 4**). The differential fractionation profiles of *Pf* Telomerase-RNP complex proteins under physiological and stress conditions hint towards their probable TERT-independent roles.

**Figure 4:**
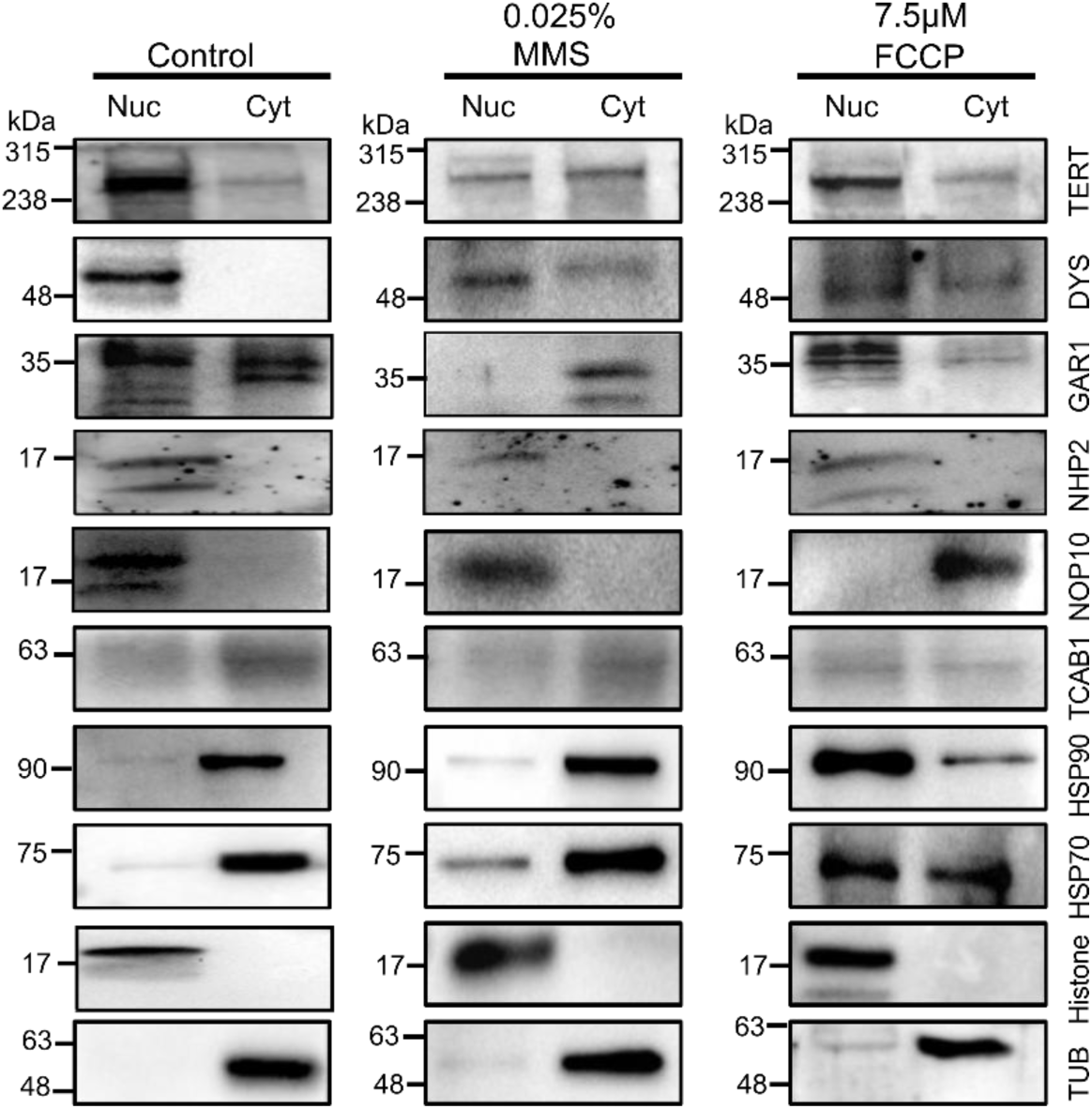
Probable TERT-independent fractionation of *Pf* Telomerase-RNP complex proteins. PF3D7 parasite culture was treated with 0.025 % MMS, 7. 5 µM FCCP at trophozoite stage (30-32 hpi) for 5.5 h. The nuclear and cytoplasmic fractions for control and treated samples were run on SDS-PAGE and western blotted with respective antibodies.

### Telomerase-RNPs as higher order complexes

Ultracentrifugation experiment was performed to investigate whether the telomerase-RNPs exist as higher order complexes in the malaria parasite. Protein complexes were separated based on their density, wherein lighter protein complexes are present in the supernatant and the heavier/denser multimeric complexes sediment into the pellet. Theoretical MW of plasmodial Telomerase-RNP complex will be ∼ 1100 kDa, and thus should sediment into the pellet fraction. Intriguingly, TERT, GAR1, NOP10 and TCAB1 were present in the both supernatant and pellet fraction, while NHP2 was predominantly in pellet fraction and Dyskerin was majorly in the supernatant (Fig 5). Probably these proteins exist as assembled and un-assembled populations due to their sub-stoichometric abundance. Upon RNaseA pre-treatment, TERT, GAR1, TCAB1 shifted to supernatant fraction. For NHP2 and NOP10, the signal was masked by non-specific reactivity with RNase in the supernatant. HSP70 was present in both fractions, with higher percentage in the supernatant, while HSP90 was obtained mainly in supernatant. RPS3 (ribosomal protein) was used as a control for ribonucleocomplexes. Under physiological condition, RPS3 exists as an unbound and ribosome-bound fraction; upon RNase treatment, RPS3 was obtained in the supernatant fraction. Tubulin was used as a cytoplasmic marker specific to the supernatant fraction.

**Figure 5:**
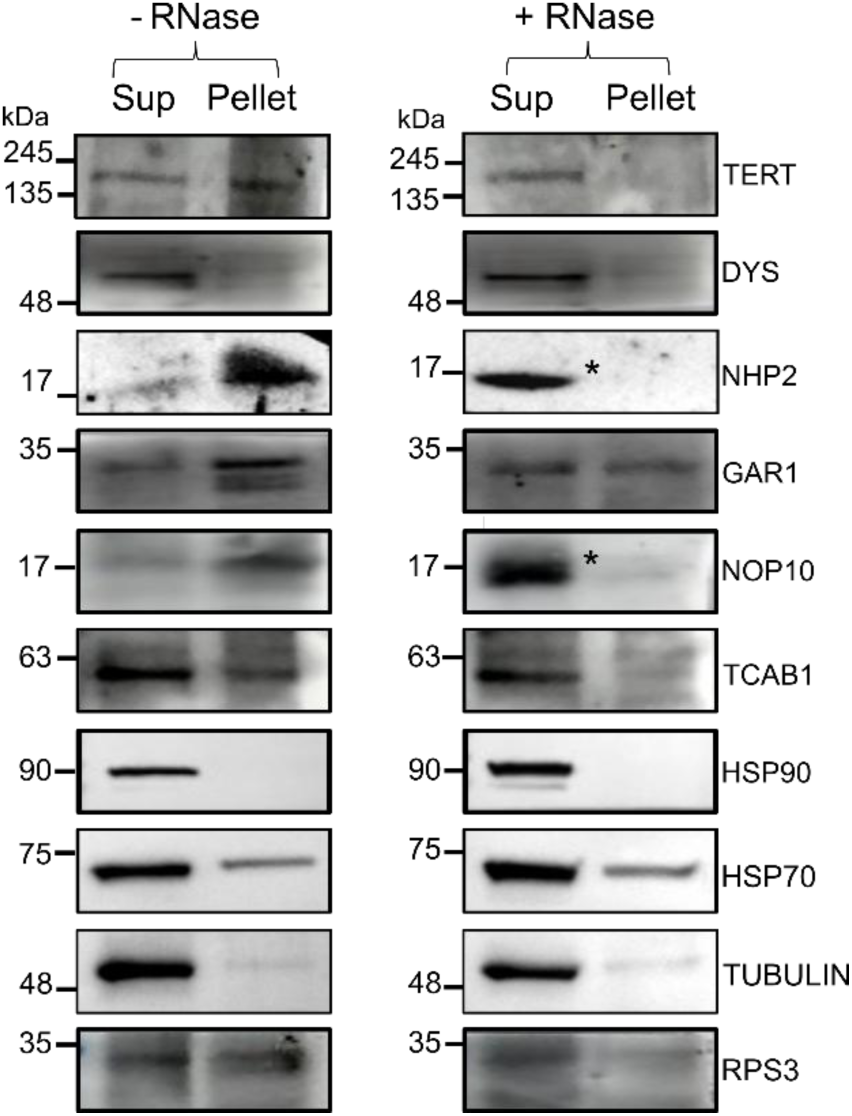
Formation of multimeric complexes by *Pf* Telomerase-RNPs. PF3D7 parasite were harvested at trophozoite stage (36-38 hpi) and protein lysate was incubated in presence or absence of 0.5 µg/µl RNase (20 °C, 15 mins). The sample was ultracentrifuged and supernatant and pellet fractions were run on SDS-PAGE. * indicates RNase band non-specifically detected by NOP10 and NHP2 antibody.

Immunoprecipitation experiments were performed to identify the interacting partners of *Pf* Telomerase. We observed significant enrichment of NOP10 in the elution fraction and no association of *Pf* TERT with GAR1. The association of Dyskerin, NHP2 and TCAB1 with *Pf* TERT could not be detected reliably. Despite limitations of this IP experiment, the data clearly suggests that these proteins may have differential binding affinity to the Telomerase-RNP complex and probably may associate transiently during the assembly of the complex. Amongst the HSPs, HSP90 formed a bonafide association with the Telomerase-RNP complex in comparison to HSP70 (**Figure S6**).

### Small molecule induced telomere damage

The evolutionary diversity of parasite telomeres and telomerase machinery can be explored for pharmacological intervention (26, 27). We performed small-molecule based parasite studies using sirtinol (a sirtuin inhibitor), quercetin (a telomerase inhibitor), and 17-AAG (an HSP90 inhibitor). The IC_50_ determined were 50 µM, 15 µM and 2.5 µM, for sirtinol, quercetin and 17-AAG, respectively. Phenotypic examination of treated parasites at ring stage (10 hpi for 18 hr) showed small-molecule induced phenotypic changes in the parasite. MMS caused enormous DNA damage and resulted in dead rings. Sirtinol inhibits Sirutin (NAD+ dependent deacetylases) (28) which play a critical role in DNA repair and metabolism. Sirtinol-treated parasites showed defects in parasite progression from ring to trophozoite stage. Quercetin and 17-AAG treatment resulted in formation of degenerated trophozoites. In addition to morphological changes, we checked their effect on telomere damage. PCR based amplification to examine telomeric length alongwith amplification of subtelomeric gene and actin as controls. MMS caused extreme loss of telomere intensity and length pointing towards the extensive DNA damage, other small molecules showed reduction in telomere intensity compared to control (**Figure 6**). These genotoxic and proteotoxic stress-inducing small molecules also affected *Pf* TERT levels, with MMS and sirtinol resulting in significant loss of *Pf* TERT expression due to induction of DNA damage (**Figure S7**).

**Figure 6:**
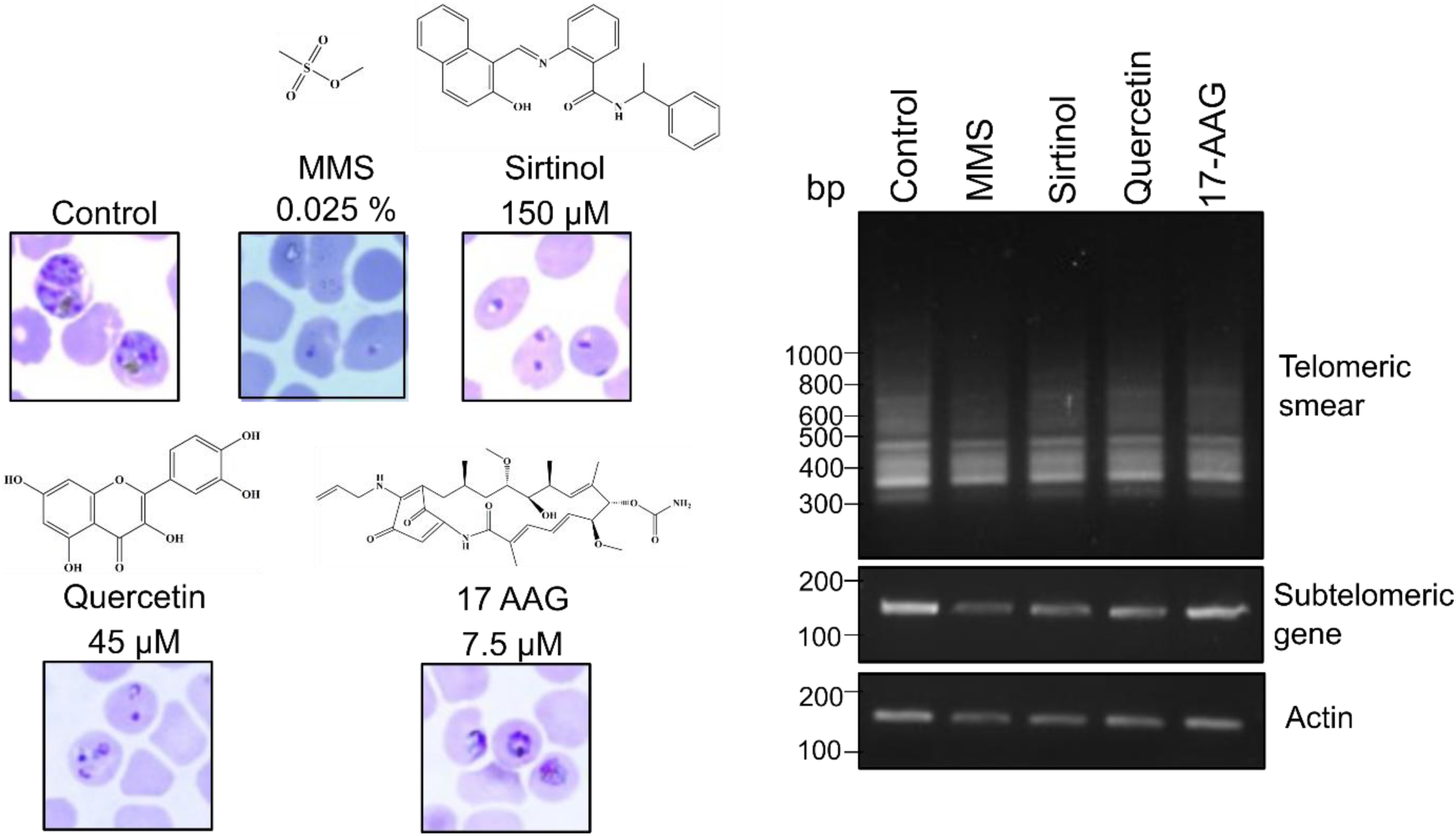
Small-molecule induced telomeric damage. Giemsa images of PF3D7 infected RBCs showing the morphological effect upon exposure of different small-molecules (treatment at ring stage for 18 h at 3× IC_50_). PCR-based estimation of telomeric length using forward primer from a sub-telomeric gene (PF3D7_1480100) and reverse primer complementary to the G-rich telomeric overhang. Amplification of actin and sub-telomeric gene (PF3D7_1480100) were used as controls.

## Discussion

The unique Telomerase-RNP complex in the human malaria parasite remains under-explored compared to other eukaryotic systems. Malaria causing species have a striking contrast in their telomere length and TERT size, with *Pf* TERT being the largest eukaryotic telomerase (Table S1). Till date, literature studies have been limited to parasite’s TERT essentiality, expression and localization (11, 15), or long read sequencing and assembly of polymorphic telomeric/subtelomeric regions (29). But these findings fall short of elucidating *Pf* Telomerase-RNP composition and its regulation. So far, are no reports on plasmodial Telomerase-structural RNA and its interacting proteins (GAR1, NOP10, DYS, NHP2, TCAB1) that assemble into a functional RNP complex to maintain the telomere length.

The substantial increase in size of *Pf* Telomerase and its non-coding RNA presents significant challenge for detailed structural characterization. The cryo-EM study of the human telomerase-RNP complex has shown that the RNA component is organized into a well-defined 5′ catalytic region, which includes the template, pseudoknot, and CR4/5 domains. These elements provide the distinct scaffolds for assembly for TERT. The 3’-end of human TR, known as the H/ACA domain adopts a double hairpin structure, with each hairpin stabilized by two molecules of Dyskerin, NOP10, NHP2, and GAR1. TCAB1 binds specifically to the P8 stem loop within the H/ACA domain, contributing to RNP stability and localization (16-18). CryoEM studies on yeast telomerase revealed Y-shaped structure, wherein its telomerase-RNA (1157 nt) forms a central core for assembly of three ever shorter telomere proteins (Ests 1-3) (30).

### Probable TERT independent roles of TR

Telomerase-RNA (TR) exhibits sequence variability across different evolutionary lineages (31). There is a growing interest in its structural versatility, which may allow TR to perform auxiliary roles in cellular homeostasis, metabolism, intracellular trafficking and stress response. A recent study on hTR used chemical probing (DMS-MaPseq) and ensemble deconvolution to understand the conformational heterogeneity and kinetic traps during its folding (19). Using chemical probing, *Trypanosoma brucei* telomerase RNA (1000 nt) architecture was shown to contain domains defining TR template, template boundary element, proximal helix and Helix IV (eCR4-CR5) (32, 33). These observations cannot be extrapolated to *Pf* TR (2148 nt) due to length and sequence diversity. Differential expression across the intraerythrocytic stages (Fig S3) and variability in region-specific amplification (Fig 2, Fig S4A) and RNA sequencing reads (PlasmoDB, Fig S4B), further support the possibility of stage-specific transcript processing or structural heterogeneity. Similar to telomerase RNAs in other eukaryotes, *Pf* TR may undergo complex maturation to generate functional RNA species; however, the highly A/T-rich parasite genome raises the possibility of species-specific processing mechanisms.

Our understanding of how *Pf* TR scaffolds TERT and other RNPs to form a catalytically active Telomerase-RNP complex is lacking, and further research is needed to address this gap in knowledge. In humans, distinct markers for Cajal-body or dense fibrillar component (DFC) are identified wherein TR folds, interacts with H/ACA interacting proteins and TERT (34). Like hTERT, *Pf* TERT shows a discrete nuclear periphery clustering rather than being diffusely nucleoplasmic **(Figure 1)**. The clustering of telomeres towards the nuclear periphery is an evolutionarily conserved phenomenon that enables better recruitment of telomerase-RNP and other regulatory complexes like shelterin for transcriptional silencing, heterochromatin formation or recombination (35).

### TERT interaction with RNP complex proteins

Literature evidence hint towards probable telomerase-independent roles of RNP complex proteins with their common role in pseudouridylation of rRNA for regulating ribosome biosynthesis. Other examples of their TERT-independent roles are (a) Dyskerin interacts with non H/ACA lncRNA to coordinate cellular survival and proliferation, (b) GAR1 participates in the dynamic assembly of phase-separated nucleolar compartments, regulating ribonucleoprotein trafficking and stability, and (c) these RNPs have been linked to ribosome-related stress signaling through p53 activation. Furthermore, their interaction with TERT may be spatially and temporally regulated. In human, it is shown that Dyskerin pre-assembles with NOP10 and NHP2 prior to encountering TR, thereby acts as a physical barrier to stop the transcription of telomerase-RNA. While, GAR1 associates with the telomerase complex in the final step of processing (24, 34, 36, 37).

The components of the Telomerase-RNP complex undergo different post-translational modifications, which may affect their stability, localization and assembly. For instance, phosphorylation and ubiquitination in TERT affects its enzymatic activity, stability and nuclear-cytoplasmic shuttling (38). As there are no reports on the *Pf* Telomerase-RNP complex, we performed comprehensive experiments on their expression and subcellular localization. TR and TR-interacting proteins are expressed in non-stoichiometric ratios to TERT (**Figures S3-S5**), hinting towards potential TERT-independent multivalent interactions such as serving as structural scaffold for other RNA binding proteins or regulating expression of genes by binding to extra-telomeric sites which may influence the cell survival and growth. We also observed differential expression and fractionation of the components of the telomerase-RNP complex (Dys, GAR1, NHP2, NOP10) under physiological and stress conditions (**Figures 3-4**). *Pf* TERT was localized primarily in nuclear fraction under physiological and stress conditions. However, GAR1 and NOP10 showed distinct nuclear to cytoplasmic trafficking during DNA damage and mitochondrial stress, respectively. We do not understand the regulatory mechanisms governing TERT-independent translocation of GAR1 and NOP10. In our immunoprecipitation experiment, GAR1 was not enriched in the eluate fraction. In human, GAR1 associates with the human telomerase complex in the final step of processing (24). GAR1 is glycine-arginine rich protein that is high-affinity stoichiometric partner of snoRNAs, and thus may have lower binding affinity to TERT (39). LC-MS based study on *Tb* Telomerase interactors identified damage specific DNA binding protein (DDB1), splicing factor 3B (SF3B1), anaphase-promoting complex (APC) and HSPs (40).

### Open questions for plasmodial Telomerase-RNP complex

The dynamics of Telomerase-RNP complex plays a critical role in the human malaria parasite as it undergoes transition from uninucleated to multinucleated stage in its evolutionary distinct hosts (human and mosquito) to complete the asexual and sexual phase of its lifecycle. As this area is underexplored in the malaria parasite, the open questions are (a) transcriptional and translational regulation mechanisms of *Pf* TERT and RNP proteins; (b) what are the post-translational modifications of *Pf* TERT and the modulators of its catalytic activity; (c) how is *Pf* TERT recruited to telomeric ends and/or to non-telomeric DNA to execute extra-telomeric roles; and (d) How does structural dynamics and conformational ensembles of *Pf* TR influence its non-canonical interactions across parasite’s lifecycle stages (41). Exploration of these questions will not only strengthen our understanding of telomere-telomerase homeostasis, but also give hint into the alternative sites of intervention. For instance, unlike humans, parasite telomeres are shorter and G-richness is predominant in telomeric and sub-telomeric ends (42-44). Targeting telomeric G-quadruplexes with selective ligands can induce selective telomeric dysregulation in the parasite (26). Another approach is developing telomerase inhibitors targeting the reverse transcriptase (RT) domain of *Pf* TERT. Low sequence and structural similarity of *Pf* TERT with human, and its constitutive expression throughout its lifecycle as opposed to cell-specific expression of *h*TERT, offers a promising opportunity for selective targeting. Additional strategy can involve structure-based discovery and validation of small-molecule modulators or antisense oligonucleotide for targeting *Pf* TR.

## Conclusion

This study provides the first comprehensive characterization of the *P. falciparum* Telomerase-ribonucleoprotein complex, highlighting its evolutionary divergence and dynamic regulation during parasite development and cellular stress. *Pf* TERT and its structural-RNA are expressed at non-stoichiometric cellular ratios with its RNP components. This suggests that the parasite has evolved a framework to meet species-specific requirements for their biogenesis, assembly to coordinate canonical (telomere maintenance, genome stability) and non-canonical roles (antigenic variation, rRNA biogenesis, stress response). Future structural and functional characterization of *Pf* TERT and its modulators may help in understanding emergence of drug-resistant persistors or quiescent subpopulations and how they bypass replicative senescence. Additionally, the striking contrast between telomere-telomerase homeostasis between human and parasite can be used for rational design of intervention strategies exploring ASOs and small-molecule inhibitor based approaches.

## Material and Methods

### Generation of constructs

*P. falciparum* genes (PF3D7_1309500, PF3D7_1123900, PF3D7_1314200) were cloned into different vector backbone (**Table S5**). Genes were amplified using Taq DNA polymerase (M829, Promega) from *P. falciparum* DNA or cDNA. The PCR product was eluted and ligated into the respective expression vector using restriction enzymes. The ligated product was transformed into *E.coli* DH5α (NEB, C29871) cells. The transformed colonies were screened and clones were confirmed by restriction digestion and sequencing. The cloning details for the respective vectors are provided in the Supplementary Information.

### STORM microscopy

Unsynchronised PF3D7 culture was maintained at 6-8 % parasitemia and 2% HCT. 30 µl of infected RBC was washed in 1× PBS thrice at 4 ⁰C at 2000 rpm for 5 mins. 500 µl of PFA (4 %) and glutaraldehyde (0.0075%) solution was added to the pack. The solution was incubated for 30 mins on RT on rotation then placed on ice for 30 more minutes till the pellet settles down. 50 µl of the pack was taken and added to the well of the quad dish. The smear was air dried in the dark. 200 µl of 30 mM glycine was added to the well for 10 mins and removed followed by one wash of 1×PBS. Then 200 µl of 0.1% triton was added on the smear. Incubated for 10 mins and then washed with 1× PBS thrice. Blocking was done using 5% BSA for 1 h at RT on rocking. BSA was removed and the primary *Pf* telomerase antibody was added to the smear and was incubated overnight at 4 ⁰C on rocking. The smear was then washed twice with 1× PBS and then incubated with secondary antibody Alexa Fluor^TM^ 647 (A-21245, Invitrogen) for 1 h at RT. Later, the smear was washed again with 1×PBS twice. Sytox dye (Invitrogen, S7020) was added and incubated for 10 mins at RT on slow rocking. The smear was then finally washed again and the well was filled with 300 µl of STORM buffer. The sample was then visualised under a custom-built setup with a Nikon Ti2E microscope body with a 647 nm laser (MPB communications). Upon excitation, the emitted light was collected by Nikon 100x oil immersion TIRF-SR objective (NA 1.49) and laser quad band set with emission filter (TRF89902-EMET-405/488/561/647 nm laser Quad Band Set for TIRF application; Chroma). EM-CCD camera was used for capturing images and hence, image localization, at an exposure time of 20 milliseconds per frame for 40,000 frames for each STORM image. Reconstruction of the image was done using a custom-written software (Insight3, provided by B. Huang, University of California, San Francisco, CA, USA).

### Expression of Telomerase-RNP proteins

*P. falciparum* (PF3D7) parasites were treated with either 0.025 % MMS (129925, Sigma,) or 7.5 μM FCCP (C2920, Sigma) at early trophozoite stage (30 hpi) for 5.5 h. Parasites were harvested at the trophozoite stage (35-36 hpi), and infected red blood cells (RBCs) were lysed using 0.2 % saponin in 1× PBS. The lysate was centrifuged at 7,000 g for 5 mins, and the resulting parasite pellet was washed thrice with 1× PBS until a clean, dark parasite pellet was obtained. The obtained parasite pellet approximately 30-40 μL of parasite pellet was resuspended in 500 μL RIPA buffer with 1 mM PMSF and 1× PIC (30 mins, 4 °C) with vortexing at 15 mins, followed by spin at 11000g (30 mins, 4 °C). the clarified lysate concentration is estimated through BCA and 70 µg of lysate is loaded for SDS-PAGE and western blotting. The nitrocellulose membranes were probed with in-house antibodies against *Pf* Telomerase, *Pf* NHP2, *Pf* GAR1, and *Pf* HSP70, as well as commercially available antibodies against NOP10 (EPR8857, Abcam), HSP90 (H1775, Sigma), and Dyskerin (SC-373956, Santa Cruz). HRP-conjugated secondary antibodies, including anti-rabbit (AP132A, Sigma), anti-rat (A5795, Sigma), and anti-mouse (A9044, Sigma), were used for detection.

For cellular fractionation experiment for separation of nuclear and cytosol, the parasite pellet was dissolved in lysis buffer (20 mM HEPES, 10 mM KCl, 1 mM EDTA, 1 mM DTT, 1 % Triton X-100, and 1× PIC). The suspension was incubated on ice for 30 min with intermittent vortexing every 5 min and subsequently centrifuged at 2,500 g for 10 min at 4°C. The supernatant was collected and designated as the cytoplasmic fraction. To remove residual cytoplasmic contamination, the pellet was washed with 1 mL of the same lysis buffer, incubated on ice for 10 min, vortexed, and centrifuged. This washing step was repeated three times to obtain a clean nuclear pellet. The resulting nuclear pellet was then resuspended in 500 μL of 1× PBS. Samples from both the cytoplasmic and nuclear fractions were prepared in 1× SDS sample dye, boiled for 20 min, and centrifuged at 12,000 rpm for 10 min. Equal volumes of the samples were resolved on 10 % and 12 % SDS–PAGE gels, followed by western blot analysis. It is important to note that we observed differences in the NHP2 bands in western blots in RIPA and fractionation buffer.

### Ultracentrifugation experiments

*P. falciparum* 3D7 culture was maintained at 6-8 % parasitemia at 2 % HCT in RPMI medium. The culture was harvested and the RBCs were lysed using 0.2 % saponin. Saponin lysed pellet was resuspended in lysis buffer (50 mM Tris pH 7.5, 150 mM NaCl, 1mM EDTA, 5 % glycerol, 1 % NP40, 0.1 % Na-Deoxycholate, 1× PIC, 1 mM PMSF), 7 times the volume of pellet. The suspension was plunged with 1 ml syringe 10 times and then incubated on ice for an hour. Lysate was then centrifuged at 12,000 rpm for 30 mins at 4 ⁰C. The protein concentration of the parasite lysate was determined using Takara BCA reagent (T9300A). The lysate was divided into two equal parts: one was incubated with 0.5 µg/µl RNaseA (101076, MP) at 20 **°**C for 15 min, while the other was kept on ice. Then it was transferred to ultracentrifuge tubes and centrifuged at 100,000 g at 4 ⁰C for 45 mins. Supernatant and pellet were separated. The obtained pellet was resuspended in 1×PBS with slight heating. The supernatant was concentrated using the TCA precipitation method. In supernatant, 0.02% sodium deoxycholate (D6750, Sigma) was added and incubated for 10 min on ice (vortex vigorously in between). 5 % Trichloroacetic acid (T6399, Sigma) was added to the solution and incubated on ice for 20 min (vortex for 1 min at every 5 min interval). The solution was centrifuged at 12,000 rpm for 20 min at 4 **°**C. The pellet formed was then washed with 1 ml ice cold acetone, incubated for 10 min and centrifuged at 12,000 rpm for 10 min at 4 **°**C, this step was repeated thrice. The pellet was dried at 62 **°**C in a dry bath and then it was resuspended in 1× SDS dye. Samples were analyzed through SDS-PAGE and western blotting with respective antibodies, *Pf* Telomerase, *Pf* HSP70, *Pf* NHP2, HSP90 (H1775, Sigma), Tubulin (T6074, Sigma), Histone (D1H2, Cell Signaling Technology), TCAB1 (BS-8163R, Bioss), NOP10 (EPR8857, Abcam) and RPS3 (NBP133691, Novus). Further, the blots were probed with their respective HRP-conjugated antibodies. BioRAD Chemidoc was used to develop the blots.

### Immunoprecipitation for interacting partners

PF3D7 at the early-late trophozoite (36-40 h) stage at 7-8% parasitemia (3x90 mm dishes) were harvested and lysed in lysis buffer (50 mM Tris pH 7.5, 150 mM NaCl, 1 mM EDTA, 5 % glycerol, 1 % NP40, 0.1 % Na-Deoxycholate, 1× PIC, 1 mM PMSF). Sepharose A beads (GE17-0780-01 GE Healthcare) were swelled overnight in water and equilibrated in the lysis buffer prior to experiment. 30 μL bed volume of sepharose A beads were incubated with 25 μL (15 μg) of anti-*Pf* TEN sera raised in rabbit for 3 h at 4°C on the tube rotator. Simultaneously, parasite lysate (600 μg) was incubated with 30 μL bed volume of sepharose A beads for 2 h at 4 °C on the tube rotator, to remove non-specific binding on beads. After incubation, both tubes were centrifuged at 2200 g for 5 min at 4 °C. Anti-*Pf* TEN sera bound beads were washed with 1× PBS (2200 g, 5 min at 4 °C) and incubated with pre-cleared lysate (600 μg) for 2 h at 4 °C on the tube rotator. Post incubation, the beads were washed with lysis buffer (2 times) and protein was eluted with 1× SDS dye (150 μL). Protein in input (60 μg) and eluate (25 μg) fractions was resolved through 10 % and 12 % SDS-PAGE, followed by western blotting. Blots were probed with *Pf* Telomerase, *Pf* NHP2, *Pf* GAR1, and *Pf* HSP70 antibodies, as well as with commercial antibodies like NOP10 (EPR8857, Abcam), HSP90 (H1775, Sigma) and Dyskerin (SC-373956, Santa Cruz). HRP-conjugated antibodies: anti-rabbit (AP132A, Sigma), anti-rat (A5795, Sigma) and anti-mouse (A9044, Sigma) were used along with the light-chain specific antibodies for HSP70 (anti-rabbit; 211-032-171, Jackson), Dyskerin (anti-mouse; 115-035-174, Jackson) and NHP2 (anti-rat; 112-035-175, Jackson).

### Isolation of parasite genomic DNA and Telomeric length Assay (TLA)

The genomic DNA isolated from parasite-infected RBC culture of trophozoite stage at 5-6 % parasitaemia. The culture was harvested by centrifugation at 2000 rpm for 5 min. The supernatant was discarded and the pellet was resuspended in 0.2% saponin solution (freshly prepared in 1× PBS) and incubated for 15 min on ice and further centrifuged at 7000 rpm for 5 mins at 4°C. The parasite pellet was washed with ice cold 1× PBS thrice to remove any RBC residues. The genomic DNA was isolated from the parasite by using ReliaPrep™ Blood gDNA Miniprep System (A5081, Promega) as per the manufacturer’s protocol. The genomic DNA was eluted in 50 μl nuclease free water.

For Telomeric length assay (TLA), *P falciparum* (PF3D7) was treated 0.025 % MMS (129925, Sigma), 150 µM Sirtinol (566320, Sigma), 45 µM Quercetin (Q4951, Sigma) and 7.5 µM 17-AAG (11039, Cayman) alongside an untreated control culture, for 18 h. The culture from the control dish and the treated dish was harvested followed by saponin lysis to isolate the parasite pellet. The gDNA was isolated from all the cultures using ReliaPrep™ Blood gDNA Miniprep System (A5081, Promega) as per the manufacturer’s protocol. 10 ng of gDNA from each condition was used as a template for 50 μl of PCR reaction. The forward primer (PF3D7_1480100_FP) was designed from the PF3D7_1480100 gene that is proximal to the telomeric end, while the reverse primers (PfCx1 and PfCx2) were complementary to the parasite’s G-rich telomeric overhang. Control PCR was performed using sub-telomeric gene (PF3D7_1480100) and actin. The sequences and PCR amplification cycle are given in **Table S5**.

### Isolation of parasite mRNA and cDNA synthesis

*P. falciparum* (PF3D7) were maintained at 6–8 % parasitemia and 2 % hematocrit in complete RPMI medium (RPMI 1640 supplemented with HEPES, 0.2 % sodium bicarbonate, 0.2 % D-glucose, 0.5 % Albumax, 45 mg/L hypoxanthine, 0.25 mg/L Fungizone, and 50 mg/L gentamicin) at 37 °C in a humidified CO_2_ incubator. RNA was isolated from different stages of parasite-infected RBC cultures. The culture was harvested by centrifugation at 2,000 rpm for 5 min at 4 °C. The supernatant was discarded, and the pellet was resuspended in TRIzol (9108, Takara) reagent (1:10, v/v) at RT and incubated for 10 min. The suspension was then centrifuged at 10,000 rpm for 5 min, and the supernatant was transferred to a fresh tube. Chloroform was added and mixed vigorously. The mixture was centrifuged at 10,000 rpm for 10 min, and the aqueous phase was passed through a gDNA removal column. The flow-through was mixed with an equal volume of 100 % ethanol and subsequently loaded onto an RNA purification column. The flow-through was discarded, and the column was washed with 500 μL of wash buffer, followed by DNase treatment for 15 min. The column was then washed twice with wash buffer, subjected to an empty spin, and the RNA was eluted in nuclease-free water (NFW). A total of 1 µg of RNA was used for cDNA synthesis using the Thermo Scientific Verso cDNA Synthesis Kit (AB1453). The 20 μL reaction mixture contained 1× cDNA synthesis buffer, dNTP mix, random hexamer primer, RT enhancer, Verso enzyme mix, RNA template, and NFW. The cDNA synthesis reaction was carried out under the following conditions: 25 ° C for 10 min, 42 °C for 1 h, and 95 °C for 10 min. Real-time PCR reactions were performed using SYBR Premix Ex Taq™ (RR420A, Tli RNaseH Plus, Takara). Prior to amplification, cDNA samples were diluted five fold and used as templates for quantitative PCR. Transcript levels of genes were analysed. Actin served as the internal reference gene. Relative RNA expression levels were normalised to actin. The primer sequences and the reaction cycle are given in **Table S6-7**

## Supplementary information

contains experimental details

## Acknowledgements

Parasite strain, 3D7 (MRA-151 contributed by David Walliker) was obtained from BEI resources NIAID, NIH. The Human Ethics Committee clearance number for *P. falciparum* culture in human RBCs is CDRI/IEC/2025/A1. The authors acknowledge DBT (BT/PR38444/MED/29/1499/2020) for funding. Authors acknowledge Ashan Manhas for assessing the antimalarial activity of inhibitors. Authors acknowledge CSIR-CDRI for infrastructural support. Authors thank Genomiki Pvt. Ltd., India for their help in analysing RNAseq-read coverage for PF3D7_0918500. CSIR-CDRI manuscript number is 148/2026/NK

## Declarations

### Ethics approval and consent to participate

Informed consent was obtained from healthy volunteers for blood for *P. falciparum* culture maintenance in human RBCs. This study is approved by Institutional Animal Ethics and Human Ethics Committee (CDRI/IEC/2025/A1).

### Consent for publication

Not Applicable

### Availability of data and material

All the data is available as Supporting information. Experimental material (plasmids) can be shared with scientific community.

### Authors’ contributions

Niti Kumar and Nirupa Chaurasia conceived the concept and supervised the study. Nirupa Chaurasia, Niharika Rai, Ariba Mahmood, Safia Baig and Neha Singh Chauhan performed all the parasite related experimental work. Nirupa Chaurasia, Niharika Rai, Suchismita Bhowmik performed STORM microscopy under the supervision of Nitin Mohan. Nirupa Chaurasia, Niharika Rai, Ariba Mahmood and Niti Kumar wrote the manuscript with inputs of other authors. All authors read and approved the final manuscript.

### Authors’ information

Niti Kumar acknowledges ANRF for SB/SJF/2021-22/03-G for fellowship. Nirupa Chaurasia and Ariba Mahmood acknowledges CSIR for fellowship, Niharika Rai acknowledges DBT, Richa Shukla acknowledges ICMR, Neha Singh Chauhan acknowledges MOES and Safia Baig acknowledges GATES foundation for their respective fellowship.

